# The Neuroanatomy of Speech Processing: A Large-Scale Lesion Study

**DOI:** 10.1101/2020.04.02.022822

**Authors:** Corianne Rogalsky, Alexandra Basilakos, Chris Rorden, Sara Pillay, Arianna N. LaCroix, Lynsey Keator, Soren Mickelsen, Steven W. Anderson, Tracy Love, Julius Fridriksson, Jeffrey Binder, Gregory Hickok

## Abstract

The neural basis of language has been studied for centuries, yet the networks critically involved in simply identifying or understanding a spoken word remain elusive. Several functional-anatomical models of critical neural substrates of receptive speech have been proposed, including (1) auditory-related regions in the left *mid-posterior* superior temporal lobe, (2) *motor*-related regions in the left frontal lobe (in normal and/or noisy conditions), the left *anterior* superior temporal lobe, or (4) *bilateral* mid-posterior superior temporal areas. One difficulty in comparing these models is that they often focus on different aspects of the sound-to-meaning pathway and are supported by different types of stimuli and tasks. Two auditory tasks that are typically used in separate studies—syllable discrimination and word comprehension—often yield different conclusions. We assessed syllable discrimination (words and nonwords) and word comprehension (clear speech and with a noise masker) in 158 individuals with focal brain damage: left (n=113) or right (n=19) hemisphere stroke, left (n=18) or right (n=8) anterior temporal lobectomy, and 26 neurologically-intact controls. Discrimination and comprehension tasks are doubly dissociable both behaviorally and neurologically. In support of a bilateral model, clear speech comprehension was near ceiling in 95% of left stroke cases and right temporal damage impaired syllable discrimination. Lesion-symptom mapping analyses for the syllable discrimination and noisy word comprehension tasks each implicated most of the left superior temporal gyrus (STG). Comprehension but not discrimination tasks also implicated the left pMTG, while discrimination but not comprehension tasks also implicated more dorsal sensorimotor regions in posterior perisylvian cortex.

---

The neural basis speech perception—defined here as the processes up to and including the successful activation of a phonological word form—has been a contentious topic for decades, if not centuries, and remains so today. Hickok & Poeppel, 2000, 2004, 2007 have reviewed the earlier literature and argued that task effects are a significant source of the lack of agreement: depending on what task is used, researchers can get different answers. In particular, they pointed out that syllable discrimination, judging whether two syllables are the same or different, can yield misleading results because they do not necessarily tap into all of the same processes involved in processing speech sounds under more normal, ecologically valid listening conditions, such as during comprehension. Specifically, it was argued that discrimination tasks recruit not only perceptual systems in the superior temporal gyrus, but also more dorsal, motor speech-related pathways. Comprehension tasks implicate perceptual systems in the superior temporal gyrus as well but do not engage the more dorsal stream and instead engage more ventral middle temporal regions. Despite highlighting these potential problems, discrimination tasks are still regularly employed to assess the neural basis of speech perception, which may perpetuate confusion and contraction in the field (Skipper, Devlin, & Lametti, 2017; Stokes, Venezia, & Hickok, 2019; Venezia, Saberi, Chubb, & Hickok, 2012). Aiming to test the task-dependence claim and to map the neural correlates of speech perception, we compared stimulus-matched word-picture matching comprehension tasks with “same-different” auditory syllable discrimination tasks in a large sample of unilateral stroke survivors. In what follows, we provide a brief summary of the major hypotheses regarding the neural basis of speech perception and then describe the present experiment. It is important to note that we do not include in our summary an important body of literature on more fine-grained aspects of speech processing in aphasia that has contributed knowledge of the psycholinguistic subtleties of disordered speech abilities and their implications for neurolinguistic models of speech processing. We exclude this work here because, unfortunately, neural correlates of the behavioral observations are not available beyond aphasia syndrome-level classification of cases. We encourage the reader to consult Blumstein (Blumstein, 1998) for a thorough review of this early literature.

Carl Wernicke (Wernicke, 1874) proposed that the left superior temporal lobe serves as the substrate for what we today generally call speech perception. The proposal is still viable today (Albouy, Benjamin, Morillon, & Zatorre, 2020) and is intuitively appealing in that the region is in auditory-related cortex and in the language dominant hemisphere. But the claim has been challenged from several angles, leading to at least three current competitors to Wernicke’s hypothesis.

The first challenge, based mostly on studies of phoneme perception using tasks such as syllable discrimination, disputes the view that sensory cortex alone is sufficient for perceptual processes, as putatively perceptual speech tasks have been found to invoke the involvement of various levels and subsystems of motor cortex (Pullvermuller et al. 2005, 2006; Rizzolatti & Arbib, 1998; Watkins et al. 2003; Wilson et al. 2004). This view was first proposed in the form of the motor theory of speech perception (Liberman, 1957, 1967) and indirectly linked the process to motor cortex. It was subsequently challenged by several examples of normal speech perception ability in the face of impaired or non-existent motor speech ability (Eimas et al. 1971; Kuhl & Miller, 1975; Bishop et al. 1990), but the motor theory claim was revived with the discovery of mirror neurons in macaque monkeys (Gallese et al. 1996). Consistent with this motor-centric view of speech perception, modern neuroimaging studies have shown that (i) motor speech regions indeed activate during listening to phonemes and syllables (Pulvermuller *et al*. 2006; Wilson *et al*. 2004; Buchsbaum *et al*. 2001; Hickok *et al*. 2003) and (ii) transcranial stimulation (TMS) studies have reported modest performance declines on syllable identification or discrimination tasks (D’Ausilio et al., 2009; Meister, Wilson, Deblieck, Wu, & Iacoboni, 2007) see (Stokes et al., 2019) for a detailed evaluation of effect size). However, strong versions of the motor centric hypothesis remain improbable for reasons given above (Hickok, 2014). A more moderate proposal in this vein is that the motor system augments speech perception only under noisy listening conditions, as broadly suggested by evidence that people with Broca’s aphasia have difficulty processing speech under degraded listening conditions (Moineau, Dronkers, & Bates, 2005; Utman, Blumstein, & Sullivan, 2001; Wilson, 2009). However, this hypothesis is complicated by the fact that lesions associated with Broca’s aphasia also often involve damage to the superior temporal lobe (Fridriksson, Fillmore, Guo, & Rorden, 2015).

A second challenge to Wernicke’s model came from neuroimaging studies at the beginning of this century showing a left dominant and specifically *anterior* superior temporal region that appeared to respond selectively to intelligible speech (Scott *et al*. 2000). Although this anterior superior temporal region has been proposed to be critical for “speech-specific processing” more broadly, the stimuli in these studies have typically been sentences (Scott et al. 2000; Evans et al. 2014; Narain et al. 2003), raising the possibility that the activation in these regions is at a post- phonological stage of processing (Rogalsky et al. 2011a; Brennan & Pylkkanen, 2012). Further, subsequent higher-powered replications of the original report show a bilateral pattern of activation involving both anterior and posterior temporal areas (McGettigan et al., 2012; Okada et al., 2010). Nonetheless, this work highlights the potential role of the anterior temporal lobe in addition to more traditional posterior temporal areas.

Finally, a third challenge came from the study of people with language disorders (aphasia) following brain injury: it was found that left hemisphere damage generally produced only mild deficits in speech perception (as measured by word to picture matching tasks with phonological foils), whereas bilateral superior temporal lesions could produce profound deficits even when pure tone hearing thresholds are normal, i.e., “word deafness” (Buchman, Garron, Trost- Cardamone, Wichter, & Schwartz, 1986; Miceli, Gainotti, Caltagirone, & Masullo, 1980; Poeppel, 2001; Henshen, 1926). In addition, studies of the isolated right hemisphere in callosotomy patients or during Wada procedures found that the right hemisphere exhibited good comprehension of most auditory words and simple phrases (Zaidel, 1985; Gazzaniga & Sperry, 1967; (Hickok et al., 2008). These findings led to the proposal that speech perception can be mediated in the superior mid to posterior temporal lobes *bilaterally* (Gazzaniga & Sperry, 1967; Buchman et al. 1986; Zaidel, 1985; Bachman & Albert, 1988; Goodglass, 1993; Hickok & Poeppel 2000, 2004, 2007; Hickok et al. 2008), which would explain the substantial effects of bilateral damage, while individuals with unilateral damage (to either hemisphere) are typically only mildly impaired on auditory word comprehension tasks. Of course, the bilateral claim is consistent with a large body of functional activation studies showing bilateral superior temporal lobe involvement in speech perception (Binder et al., 2000; DeWitt & Rauschecker, 2012, 2016; Hickok & Poeppel, 2007; Leonard, Baud, Sjerps, & Chang, 2016; Moses, Mesgarani, Leonard, & Chang, 2016; Myers, 2007; Okada & Hickok, 2006; Okada et al., 2010; Overath, McDermott, Zarate, & Poeppel, 2015).

Much of the modern research on this topic has been carried out using neuroimaging methods, which cannot assay the causal involvement of activated networks. There have been attempts to assess the competing proposals more directly using neuropsychological (Bak & Hodges, 2004) or, as noted above, neural interference methods (TMS) (D’Ausilio *et al*. 2009; Meister *et al*. 2007; Mottonen & Watkins, 2009). However, questions have been raised about the generalizability of these results to natural receptive speech processing, particularly for the TMS studies, because the reported effects on perceptual accuracy are small (estimated to amount to a decrement of only about 1-2 dB (Stokes et al., 2019), are only found for syllable discrimination or identification tasks not comprehension (Schomers, Kirilina, Weigand, Bajbouj, & Pulvermuller, 2014), and are evident only in near-threshold listening conditions (see (Hickok, 2014; Stokes et al., 2019) for discussion). Thus, despite more than a century of study, the neural basis of speech perception remains controversial.

To summarize, we have outlined the following four hypotheses regarding the neural basis of speech perception, again, defined as the processes up to and including the successful activation of a phonological word form:

1. Classical model—speech perception is uniquely supported by the left mid-to-posterior superior temporal lobe.
2. Motor model—speech perception is dependent on (strong version) or augmented by (weak version) one or more motor sub-systems involved in speech production. Augmentation for speech perception under the weak model is typically assumed to hold for noisy or otherwise effortful listening conditions.
3. Left anterior temporal model—speech perception is dependent on the left anterior superior temporal lobe.
4. Bilateral temporal model—speech perception is bilaterally organized in the mid-to-posterior superior temporal lobe.

We have also outlined an additional claim that discrimination and comprehension tasks implicate shared networks in the superior temporal gyrus but then engage different dorsal (discrimination) versus ventral (comprehension) pathways beyond this region. Thus,

5. Task dependent claim—speech processing as measured by discrimination tasks will implicate the mid-to-posterior superior temporal gyrus and more dorsal regions, while speech processing as measured by comprehension will implicate the mid-to-posterior superior temporal gyrus and more ventral regions.

To test these hypotheses, we conducted a large-scale lesion study employing two auditory word comprehension tasks and two syllable discrimination tasks. Both of the auditory word comprehension tasks involved matching a spoken word to a picture in an array of four where one picture was the target and the other three were phonological, semantic, or unrelated distractors.

In one version of the task, speech was presented in a quiet background (clear speech) and in the other version, speech was presented in white noise. The speech-in-noise task served two purposes, one to test claims about the role of the motor system in processing noisy speech and second, to ensure that the distribution of performance on word comprehension is below ceiling, thus enabling lesion mapping.

The syllable discrimination tasks involved the presentation of pairs of monosyllabic speech stimuli with minimally contrasting onsets and a 1 second inter-stimulus interval. The listener was asked to decide whether the two stimuli were the same or different. One version of the task involved real words and the other involved nonwords. Despite the intuitive assumption that both discrimination and auditory word comprehension tasks measure an ability to extract sound-based phonological information from a speech signal, these two tasks are reported to be doubly- dissociable (Hickok & Poeppel, 2004; Miceli, Gainotti, Caltagirone, & Masullo, 1980).

Particularly paradoxical are cases in which the ability to discriminate pairs of syllables (e.g., *da* from *ba*) is impaired while the ability to differentiate words with these same sounds (e.g., *doll* versus *ball*) is spared in comprehension tasks. This has confounded models of the neural basis of speech perception and recognition for decades because one can reach different conclusions depending on which task is used (Hickok, 2009, 2014; Hickok & Poeppel, 2000). We sought to resolve this issue in the present study by employing both tasks in a large sample. We studied people with either left or right hemisphere chronic stroke, left or right anterior temporal lobectomy, and neurologically intact controls. For our largest group (the left hemisphere stroke group) we use lesion-behavior mapping to identify brain regions responsible for receptive speech deficits.

## Predictions

Based on their evaluation of previously published studies, Hickok & Poeppel (2000, 2004, 2007) predict that word comprehension and syllable discrimination tasks are doubly dissociable both behaviorally and neurally, even when the syllables in the discrimination task are words. This would argue strongly that discrimination tasks add some process to speech perception that is not necessarily used during normal word comprehension (see (Burton, Small, & Blumstein, 2000). Assuming that researchers are aiming to understand how speech is processed “in the wild” (this is indeed our own goal) and assuming that the word comprehension task is a closer approximation to ecological validity, such a result would seriously question the validity of discrimination tasks in studies of speech perception. Turning to the neuroanatomical models, the *classical model* predicts that damage to the posterior superior temporal lobe should yield substantial deficits on both tasks. *The motor models* predict that damage to frontal, motor-related regions—which may include primary motor cortex (D’Ausilio et al., 2009), premotor cortex (Meister et al., 2007), inferior frontal gyrus (Burton et al., 2000; Mottonen & Watkins, 2009), or subparts thereof—should cause speech perception deficits. In the present study we sample all of these regions and so will be in position to test each of these possibilities. The strong variant predicts deficits on all tasks, whereas the weak variant predicts impairment on tasks involving speech in noise. The *left anterior temporal model* predicts that unilateral mid-to-anterior superior temporal lobe damage should substantially interfere with word comprehension via disruption to systems critical for recovering intelligibility. It is less clear what predictions this model has for nonword discrimination. The *bilateral temporal model* generally predicts minimal impairment on speech perception tasks, perhaps of any type, following unilateral damage. The *task-dependent claim* predicts overlapping lesion correlates for syllable discrimination and word comprehension tasks in the superior temporal gyrus, with differences emerging in dorsal regions (discrimination only) and ventral regions (comprehension only).

## Methods

### Participants

Consent from all participants (described below) was obtained according to the Declaration of Helsinki and the experiments were approved by the Institutional Review Boards of all the institutions at which the work was performed.

#### Lesion Participants

The following groups of participants were examined: 113 individuals who had previously experienced a left hemisphere stroke (Figure 1), 19 individuals who had previously experienced a right hemisphere stroke (Figure 2 A,B), 18 individuals who had previously underwent a unilateral left temporal lobectomy, and 8 individuals who had previously underwent a unilateral right temporal lobectomy (Figure 2 C,D). See Table 1 for demographic information of each group. The left hemisphere stroke participants were tested at six institutions: University of Iowa (n=40), University of South Carolina (n=32), Medical College of Wisconsin (n=24), San Diego State University (n=9), Arizona State University (n=7) and University of California, Irvine (n=1). The right hemisphere stroke participants were tested at University of Iowa (n=17), University of South Carolina (n=1) and Arizona State University (n=1). All lobectomy participants were tested at the University of Iowa. Participants were included in the present study based on the following criteria: (i) a chronic focal (6 months or more post-onset) lesion due to a single stroke in either the left or right cerebral hemisphere or a unilateral temporal lobectomy (ii) no significant anatomical abnormalities other than the signature lesion of their vascular event (or evidence of surgery for the lobectomy subjects) nor signs of multiple strokes, (iii) an absence of a history of psychological or neurological disease other than stroke (or seizure disorder for the lobectomy subjects), (iv) native English speaker, (v) right handed pre-stroke, and (v) ability to follow task instructions. The temporal lobectomy patients all had a seizure disorder that required lobectomy surgery for the treatment of their seizures^1^. All stroke participants had ischemic strokes with the exception of two participants in the left hemisphere group with hemorrhagic strokes.

**Figure 1.**
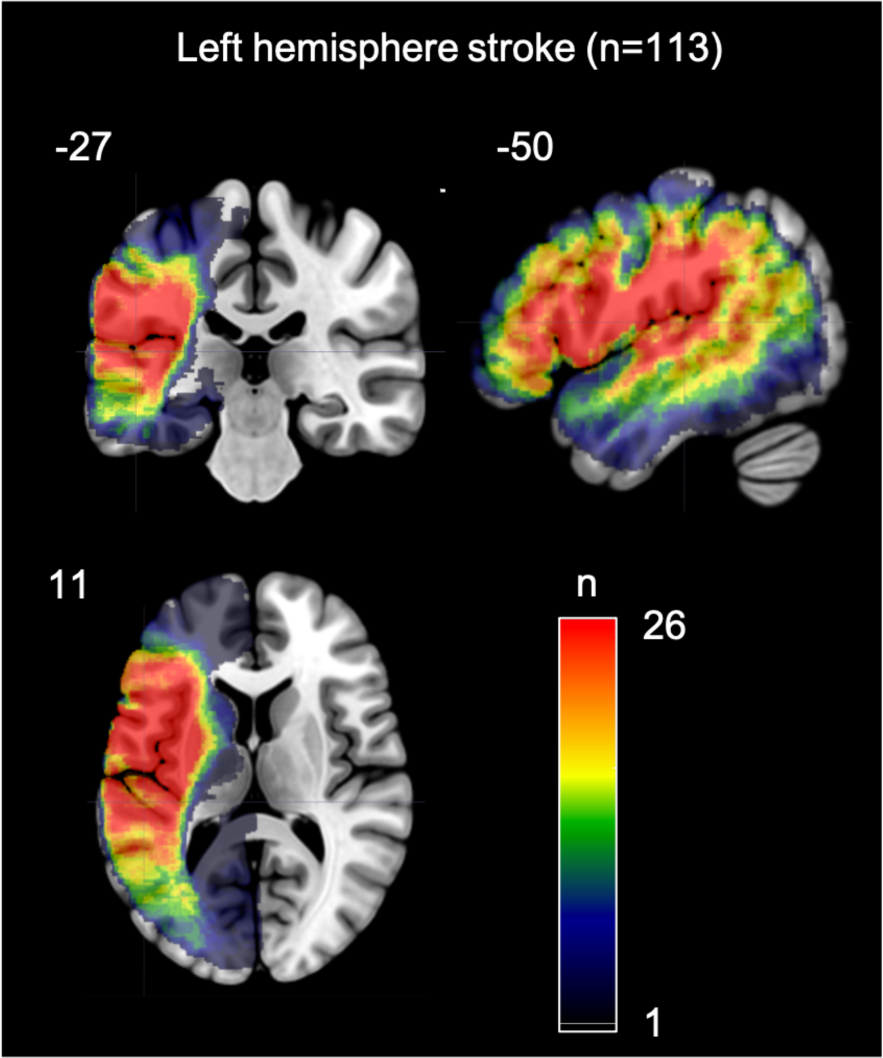
Overlap map of the areas of damage in the 113 participants with a left hemisphere stroke.

**Figure 2.**
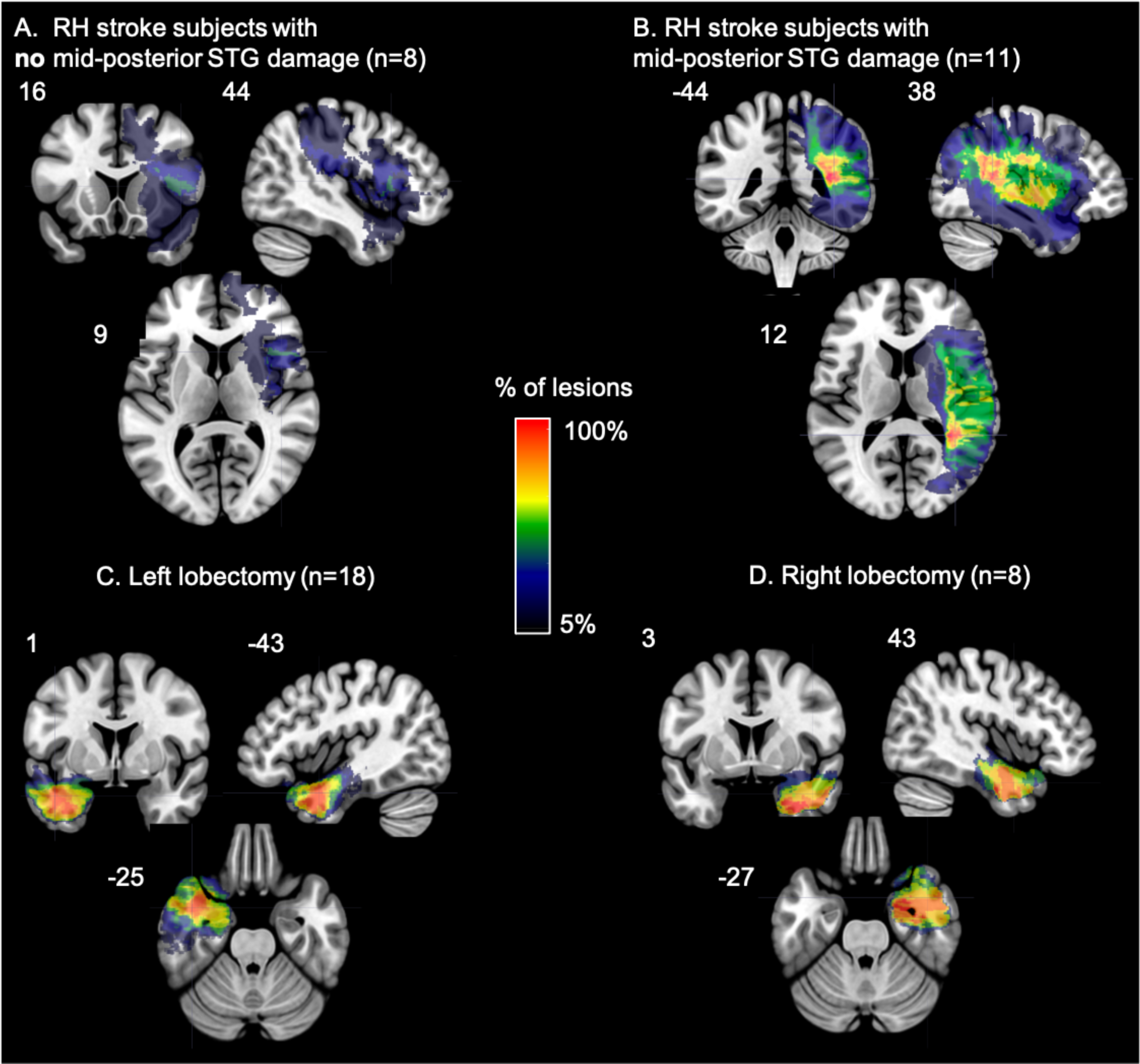
(A,B): Overlap map of the areas of damage in the participants with a right hemisphere stroke (A) without and (B) with mid-posterior STG damage. (C,D): Overlap map of the areas resected in the participants who underwent a (C) left or a (D) right temporal resection.

**Table 1.** Demographic information for each participant group. Note that “time post” refers to the number of years since a stroke for the left hemisphere and right hemisphere stroke groups, and refers to the number of years post-resection for the left lobectomy and right lobectomy groups.

|  | n | Age<br>(years) | Gender<br>(M/F) | Education<br>(years) | time post<br>(years) |
| --- | --- | --- | --- | --- | --- |
| Left Hemi. Stroke | 113 | 61 (31-86) | 67 / 46 | 15 (8-20) | 6.1 (0.5-32) |
| Right Hemi. Stroke | 19 | 52 (25-75) | 13 / 6 | 15 (10-20) | 11 (0.5-35) |
| Left Lobectomy | 18 | 48 (29-62) | 8 / 10 | 13.5 (10-16) | 4 (0.7-14) |
| Right Lobectomy | 8 | 41 (23-60) | 3 / 5 | 14 (12-18) | 9 (4-18) |
| Control | 26 | 59 (12-16) | 5 / 21 | 15 (12-16) | n/a |

Aphasia type was assessed in the stroke and lobectomy groups via site-specific protocols that included the Western Aphasia Battery, Boston Diagnostic Aphasia Examination, and clinical ratings. The left hemisphere stroke group consisted of the following aphasia diagnoses: Broca’s (n=30), anomic (n=15), conduction (n=9), residual aphasia (n=8), Wernicke’s (n=7), transcortical motor (n=5), mixed non-fluent (n=5), global (n=3), mild Broca’s / anomic (n=2), and no aphasia (n = 29). None of the participants in the lobectomy groups or in the right hemisphere stroke group were diagnosed with aphasia.

Given the proposed role of the right mid-posterior STG in speech perception, the right hemisphere stroke patients were divided into two groups: those with right mid-posterior STG damage (n=11) and those with no mid-posterior STG damage (n=8; max overlap is 3/8 subjects in portions of the inferior frontal gyrus, precentral gyrus and insula) (Figure 2A,B). Mid-posterior STG was defined as any gray or white damage in the STG adjacent or posterior to Heschl’s gyrus as determined by visual inspection by one of the authors (Rogalsky) with extensive training in cortical neuroanatomy. The area of maximum lesion overlap for the RH stroke participants with mid-posterior STG damage is located in white matter underlying the posterior temporal lobe more generally.

*Control Subjects.* Twenty-six adults were recruited and tested at San Diego State University. All control subjects were native English speakers, right-handed, and self-reported no history of psychological or neurological disease. See Table 1 for demographic information.

Informed consent was obtained from each participant prior to participation in the study, and all procedures were approved by the Institutional Review Boards of UC Irvine, University of South Carolina, San Diego State University, Arizona State University, Medical College of Wisconsin and University of Iowa.

Standard hearing and vision screenings were performed during the recruitment of all participants to ensure adequate hearing and vision abilities to complete the experimental tasks and aphasia assessments. The exact hearing and vision screenings varied across sites, but also all participants were provided with several practice trials for each task during which presentation volume could be adjusted to each participant’s preferences.

### Materials

The present study focuses on two types of tasks: auditory word comprehension and syllable discrimination. The tasks were administered as part of an extensive psycholinguistic test battery to assess receptive and productive speech abilities. Within the battery, individual tests themselves were presented in a non-fixed pseudorandom order; The order and proximity within the test battery of the tasks described below was random across subjects. Items within each test described below were presented in a fixed random order at all testing sites, except for Medical College Wisconsin (n=24), where items were presented in a random order for each subject.

### Auditory Comprehension Measures

#### Auditory word comprehension in a clear background

A four alternative forced choice word-to-picture matching task previously used by Rogalsky et al. (2011b) and similar to Baker, Blumstein, and Goodglass’ (1981; Blumstein *et al*. 1977) paradigm was administered. The task contained 20 trials. In each trial an auditory word was presented one second after the appearance of a picture array depicting four objects on a computer screen. The words were recorded by a male native speaker of American English. The participant was instructed to point to the picture that matched the auditorily-presented word. The picture array contained the target and three distractors. Ten of the 20 trials included a one syllable target word and its matching picture, as well as a picture representing the phonemic distractor that was a minimal pair to the target (4 place of articulation contrasts, 4 voicing contrasts, and 2 manner contrasts), a semantic distractor, and an unrelated distractor, which was semantically related to the phonological distractor. For example, the target *goat* was presented with distractor images of a coat, a pig, and a shoe. The auditory stimuli were counterbalanced such that each picture in each array was presented once as the target. All of the words presented have an estimated average age of acquisition under six years old (range: 2.6- 5.7 years old; Kuperman et al. 2012) and ranged in frequency from 1.33 to 86.86 per million words (Brysbaert & New, 2009). The frequency of each error type was calculated, as well as proportion correct for each subject’s overall performance in each task.

### Auditory word comprehension in noise

The auditory comprehension in noise task contains the same word and picture array pairings as the auditory comprehension word-picture matching task described above, except that the auditory stimuli were presented in 14 dB Gaussian white noise. The noise began one second before the onset of the word and ended one second after the end of the word.

### Syllable Discrimination Measures

We employed two syllable discrimination measures: a real word discrimination task and a nonword discrimination task. These two syllable discrimination tasks were modeled on previous discrimination tasks used with individuals with aphasia (Baker et al. 1981; Blumstein et al. 1977; Caplan & Waters, 1995). Each of these discrimination tasks contained 40 trials. The real word discrimination task contained pairs of one syllable real words and the nonword discrimination task contained one-syllable nonwords, with neighborhood density and phonotactic probability between the real words and nonwords matched. The Irvine Phonotactic Online Dictionary (Vaden et al. 2009) was used to select the real words and nonwords, and to calculate neighborhood density and phonotactic probability for each word and nonword. Each word and its corresponding nonword have the same onset consonant phoneme. Paired-samples t-tests comparing each word with its matched nonword indicate that there is no significant difference between the two stimulus types for density *t*(39) = .14, *p* = .88 or phonotactic probability *t*(39) = .18, *p* = .86. The words used in the real word discrimination task do not overlap with the words in the auditory comprehension tasks described above, but the minimal pairs in the syllable discrimination tasks are the same minimal pairs present in the auditory word comprehension tasks.

The real word and nonword discrimination tasks both were structured in the following manner: There were 40 trials in each task, and in each trial, a pair of one-syllable words (or nonwords) were presented via headphones while a fixation cross was presented on the computer screen.

Each trial presented the two words (or nonwords) in one of four arrangements: A-B, B-A, A-A, and B-B such that for half of the trials the correct answer was “same” and for the other half the correct answer was “different”. In the “same” trials, two different tokens of the same syllable were presented. The same male native English speaker of the words in the comprehension task described above recorded all of the stimuli. There was a one second interval between each word (or nonword) in a pair, and five seconds between each pair (more time between pairs was given, if necessary, for the patient’s comfort). Patients were instructed to determine if the two words (or nonwords) presented were the same or different. “Different” trials contained two syllables that differed by one feature of the onset consonant (e.g. “puff” vs “cuff” in the real word discrimination task, or “pag” vs “kag” in the nonword discrimination task). Participants made their response either by speaking their answer or by pointing to the correct answer written in large print on a paper adjacent to the computer screen. Signal detection methods were used to determine how well subjects could discriminate between the same and different pairs by calculating the measure d’ for the real word and nonword discrimination tasks, respectively (Swets, 1964).

### Neuroimaging

All stroke and lobectomy participants (n=158) underwent MRI scanning using a 3T or 1.5T MRI system at the respective testing site. T1-MRIs and T2-MRIs with 1 mm^3^ resolution were collected and used to demarcate the lesion; the lesion demarcation was conducted manually by well-trained and experienced individuals in lesion-mapping studies. The lesion maps were smoothed with a 3mm full-width half maximum Gaussian kernel to remove jagged edges associated with manual drawing. Enantiomorphic normalization (Nachev *et al*. 2008) was conducted using SPM12 and ‘in house’ Matlab scripts in the following way: a mirrored image of the T1 image (reflected around the midline) was coregistered to the native T1 image. Then, we created a chimeric image based on the native T1 image with the lesioned tissue replaced by tissue from the mirrored image (using the smoothed lesion map to modulate this blending, feathering the lesion edge). SPM12’s unified segmentation-normalization (Ashburner & Friston, 2005) was used to warp this chimeric image to standard space, with the resulting spatial transform applied to the actual T1 image as well as the lesion map. The normalized lesion map was then binarized, using a 50% probability threshold.

### Lesion-Symptom Mapping (LSM)

All of the LSM routines used here are integrated into the NiiStat toolbox for Matlab (http://www.nitrc.org/projects/niistat). The following routines were performed: Univariate analyses fitting a general linear model (GLM) were completed to identify anatomical regions in which the percentage of lesioned voxels was associated with performance on the auditory comprehension in noise, word discrimination, or nonword discrimination tasks. A GLM was fit for performance on each task within each of the 94 left hemisphere regions of interest (ROIs) defined by the John’s Hopkins University (JHU) atlas (Mori *et al*. 2008; Faria *et al*. 2012; den Ouden *et al*. 2019). An ROI approach, compared to voxel-based statistics, increases statistical power by reducing the severity of multiple comparison correction (Rorden et al. 2009).

Univariate analyses were conducted for the following variables: auditory word comprehension in noise (proportion correct), word discrimination (d’) and nonword discrimination (d’). Clear speech auditory word comprehension was not used for LSM due to a ceiling effect with insufficient variability; we discuss this further below. The behavioral data were significantly skewed (i.e. Z-Skew > 1.96) thus de-skewing was performed in NiiStat prior input into the GLM by computing the standard error of the skew and applying a square root transform to the data.

Permutation thresholding included 4,000 permutations to correct for multiple comparisons (p < 0.05 controlled for familywise error). As ROIs that are infrequently damaged will have low statistical power while increasing the number of comparisons, only ROIs where at least 11 participants (i.e. approximately 10% of the largest left hemisphere patient sample size in a task) had damage were included in the analyses.

Variability due to overall lesion size was regressed out of each LSM. Data collection site also was used as a covariate to remove variance accounted for by collection site, including potential variance due to site-specific differences in scanner properties and protocols. All reported coordinates are in MNI space. Figure 3 displays the extent of coverage of the ROIs included in the lesion-symptom mapping analyses; the coverage does not vary between the analyses.

**Figure 3.**
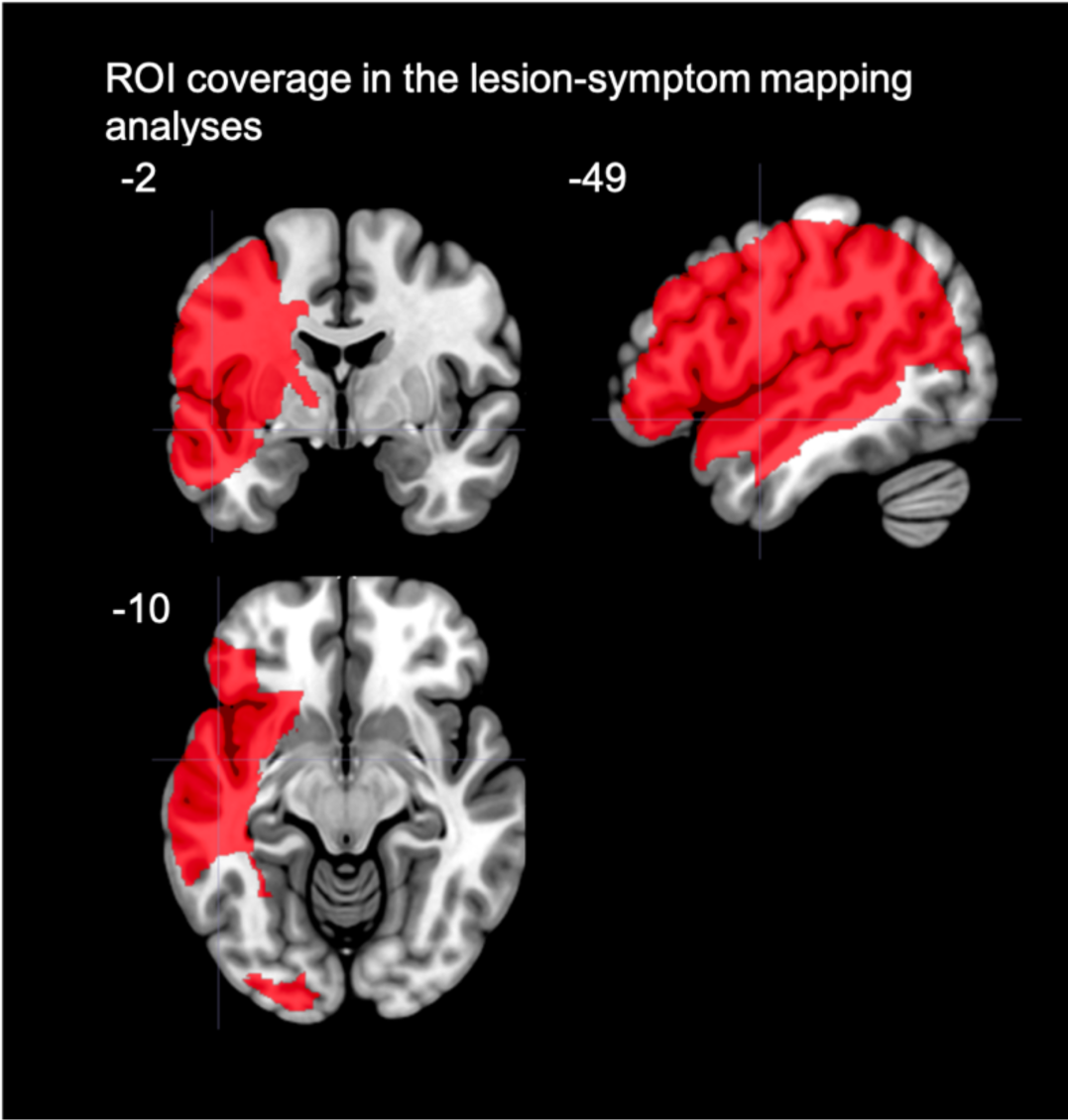
Map of the extent of coverage by the ROIs from the JHU atlas included in the lesion-symptom mapping analyses of the left hemisphere group. Note the ample coverage of the superior and middle portions of the temporal lobe, as well as premotor, inferior frontal, and primary motor cortex.

### Post-hoc comparisons of LSMs

Based on the close proximity and partial overlap of the significant ROIs identified by the LSMs described above, we decided to use conjunction and independence tests to further investigate the relationships between (1) the brain regions supporting real word discrimination versus those regions supporting nonword discrimination, and (2) the brain regions supporting nonword discrimination versus auditory comprehension in noise. The conjunction tests were computed in each ROI using an in-house Matlab script following the valid conjunction test procedure recommended by Nichols et al. (2005). Code for the conjunction analysis can be found here: https://raw.githubusercontent.com/rordenlab/spmScripts/master/nii_thresh_conjunction.m. This conjunction analysis procedure generates, in each ROI, the p-value of the z-values under the conjunction null, i.e. the union of the null hypotheses for the two tasks’ z-scores. ROIs with p < .05 indicate that damage in that ROI is associated with lower performance in both tasks. The independence tests were computed in each ROI using a custom contrast in the nii_stat toolbox (e.g. 1 -1) to determine if damage in each ROI is significantly more related to one task than the other; in other words, if an ROI’s z-score for one task is significantly different than the z-score for another task (*p*< .05, Freedman-Lane permutation as described by Winkler et al. 2014).

Due to a combination of time limitations and site-specific testing protocols, not all participants completed all speech perception tasks. Table 1 indicates the sample sizes for each task within each participant group. Within each lesion group, there are no qualitative differences in the lesion distributions or lesion sizes of the participants who did and did not complete each task.

## Results

### Overall Behavioral Results

A summary of the descriptive statistics for each group on each of the tasks is presented in Table 2.

**Table 2.** Descriptive statistics for the clear auditory word comprehension (proportion correct), auditory word comprehension in noise (proportion correct), word discrimination (d-prime values), and nonword discrimination (d-prime values) tasks.

| Clear Auditory Word Comprehension |  |  |  |  |  |
| --- | --- | --- | --- | --- | --- |
|  | Control | LH Stroke | RH Stroke | Left ATL | Right ATL |
| **n** | 21 | 109 | 19 | 18 | 8 |
| **Mean** | 1 | 0.97 | 0.99 | 1 | 1 |
| **Median** | 1 | 1 | 1 | 1 | 1 |
| **Std. Deviation** | 0 | 0.08 | 0.03 | 0 | 0 |
| **Range** | 1-1 | 0.4-1 | 0.9-1 | 1-1 | 1-1 |

| Auditory Word Comprehension in Noise |  |  |  |  |  |
| --- | --- | --- | --- | --- | --- |
|  | Control | LH Stroke | RH Stroke | Left ATL | Right ATL |
| **n** | 22 | 109 | 19 | 17 | 8 |
| **Mean** | 0.91 | 0.78 | 0.88 | 0.87 | 0.91 |
| **Median** | 0.90 | 0.80 | 0.90 | 0.85 | 0.90 |
| **Std. Deviation** | 0.05 | 0.14 | 0.05 | 0.07 | 0.05 |
| **Range** | 0.85-1 | 0.3-1 | 0.8-0.95 | 0.75-1 | 0.85-1 |

| Word Discrimination |  |  |  |  |  |
| --- | --- | --- | --- | --- | --- |
|  | Control | LH Stroke | RH Stroke | Left ATL | Right ATL |
| **n** | 25 | 78 | 18 | 17 | 8 |
| **Mean** | 5.05 | 4.20 | 4.72 | 5.02 | 5.15 |
| **Median** | 5.15 | 4.22 | 5.15 | 5.15 | 5.15 |
| **Std. Deviation** | 0.29 | 1.06 | 0.78 | 0.38 | 0 |
| **Range** | 4.22-5.15 | 1.71-5.15 | 2.68-5.15 | 3.86-5.15 | 5.15-5.15 |

| Nonword Discrimination |  |  |  |  |  |
| --- | --- | --- | --- | --- | --- |
|  | Control | LH Stroke | RH Stroke | Left ATL | Right ATL |
| **n** | 23 | 77 | 18 | 17 | 8 |
| **Mean** | 4.79 | 3.75 | 4.29 | 4.58 | 4.76 |
| **Median** | 5.15 | 4.22 | 4.22 | 5.15 | 5.15 |
| **Std. Deviation** | 0.57 | 1.27 | 1.02 | 0.86 | 0.56 |
| **Range** | 3.61-5.15 | 0-5.15 | 2.07-5.15 | 2.68-5.15 | 3.86-5.15 |

#### Auditory word comprehension tasks

all groups were at ceiling on the clear auditory word comprehension task. In the left hemisphere stroke group, the mean performance was 97% correct (Figure 4) with both the left and right anterior temporal lobectomy groups scoring at 100% correct. More variance in performance was found on the auditory word comprehension in noise task, enabling statistical analyses. A one-way ANOVA across the 5 groups (control, LH stroke, RH stroke, L lobectomy, R lobectomy) revealed a significant group effect in performance on the auditory word comprehension in noise task (F(4, 170) = 10.6, p < .001) with the LH Stroke group differing from all other groups (*p*s *≤* .002, Bonferroni correction of *α*= .05/10 =.005) and no other group differences (Figure 5, left panel).

**Figure 4.**
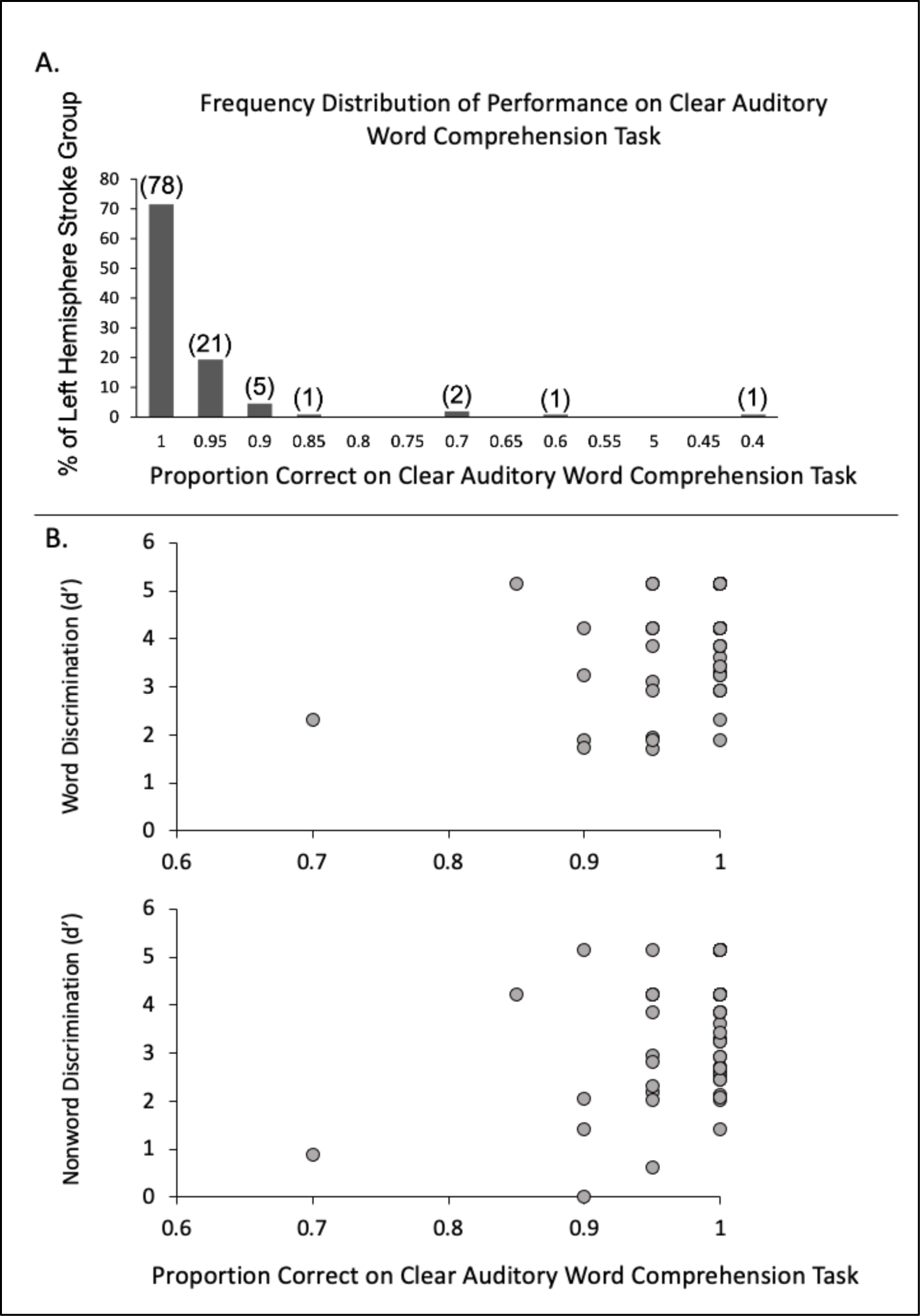
A. Distribution of performance of the left hemisphere stroke group on the clear auditory comprehension task. Numbers in parentheses indicate the actual number of left hemisphere stroke participants who performed at that accuracy level (out of a total 109 who completed the task). Note that chance performance would be .25 proportion correct (not shown). B. Distribution of performance of the left hemisphere stroke group on the clear auditory comprehension task as a function of word discrimination (top) and nonword discrimination (bottom) task performance.

**Figure 5.**
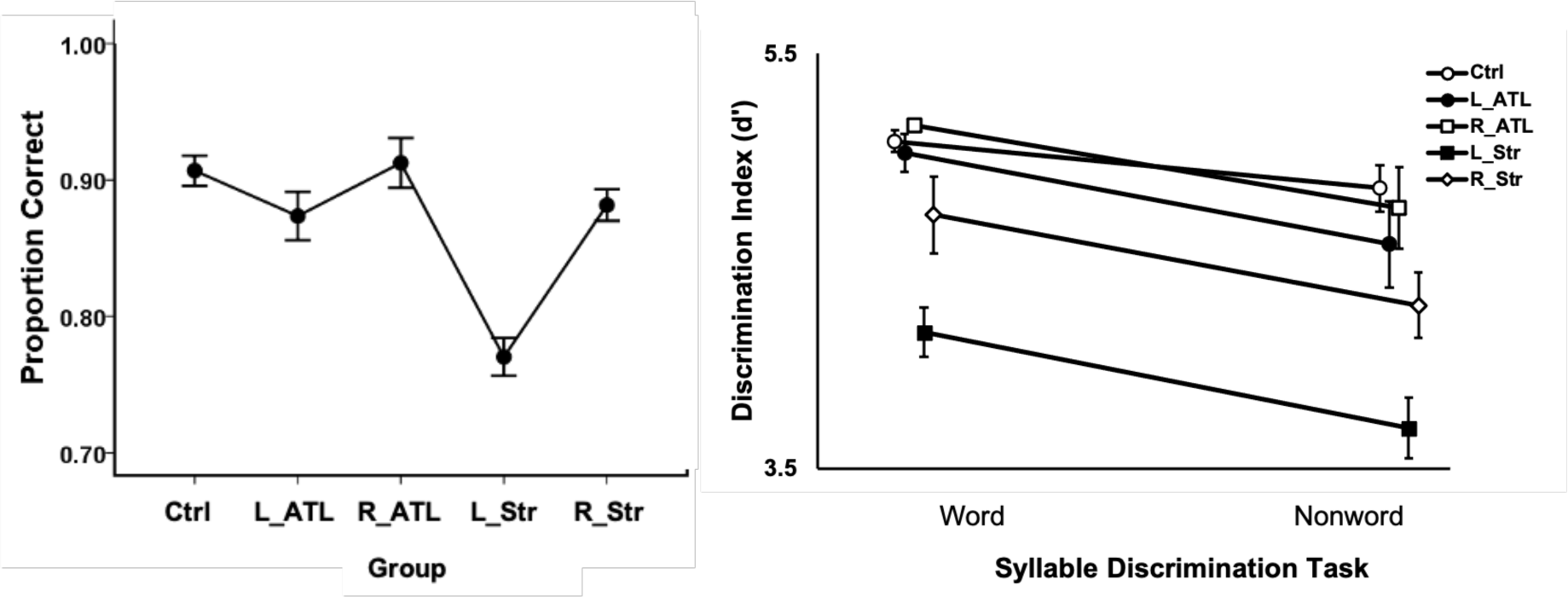
Performance on the auditory word comprehension in noise (left) and syllable discrimination tasks (right) in each participant group. Error bars represent standard error. Overlapping points in the discrimination task graph have been offset slightly to make them visible. Ctrl = control group, ATL = lobectomy, Str = stroke.

#### Syllable discrimination tasks

a 5 x 2 mixed ANOVA was computed for the performance on the two syllable discrimination tasks (i.e. word discrimination and nonword discrimination) in each of the five groups revealing significant main effects of Task (F(1,138) = 17.80, p < .001) with higher scores on the word than nonword discrimination, and Group (F(4, 138) = 9.20, p < .001). The interaction was not significant (F(4,138) = 0.38, p=.83). The LH Stroke group performed significantly worse than the control, L lobectomy, and R lobectomy groups (*p*s *≤* .002, Bonferroni correction of *α* = .05/10 =.005) with no other significant group differences. The LH stroke group numerically performed worse than the RH stroke group across both syllable discrimination tasks, but the comparison did not withstand multiple comparison correction (*p*= .015) (Figure 5, right panel).

As noted in the introduction, previous observations indicated that performance on auditory word comprehension and syllable discrimination tasks are doubly dissociable. We assessed this in our sample by categorizing performance as impaired or spared on the auditory word comprehension in noise task and the word discrimination task. “Impaired” was defined as performance greater than two standard deviations below the control mean; “spared” was defined as performance that is within or above .5 standard deviations of the control mean. Seventy-five LH stroke participants completed both of these tasks. Table 3 presents participant counts that fall into the 4 possible spared-impaired categories on the two tasks. It is apparent that these two tasks are dissociable, with 13 cases (17.3%) exhibiting impaired performance on the auditory word comprehension task but spared performance on the word discrimination task, and 10 cases (13.3%) showing the reverse pattern. There are fewer cases with spared auditory word comprehension in noise in the face of impaired word discrimination but recall that performance on *clear* word comprehension is at ceiling for the majority of LH Stroke participants. There are 59 such participants who were both at ceiling on the clear auditory word comprehension task and completed the word discrimination task; 30 (50.8%) were impaired on the latter, which was also presented as clear speech. This provides substantial evidence that these tasks are dissociable.

**Table 3.** Frequency counts of LH Stroke cases that fall into the four categories of impaired and spared performance. Impaired is defined as performance that is > 2 standard deviations below the control mean; spared is defined as performance that is within or above .5 standard deviations of the control mean.

| Auditory Word Comprehension in Noise | Word Discrimination |  |
| --- | --- | --- |
|  | Impaired | Spared |
| Impaired | 26 | 13 |
| Spared | 10 | 12 |

### Word Discrimination

Within the LH stroke group, correlation analyses performed between the auditory word comprehension in noise and word discrimination tasks revealed a statistically significant relationship (*r*(73) = .39, r^2 =^ .15, *p* <.001). However, it is evident from the scatterplot in Figure 6 that, consistent with the categorization-based analysis of task dissociability, continuous variation in performance on one task is a rather poor predictor of variation on the other, accounting for only 15% of the variance. A partial correlation between the auditory word comprehension in noise and word discrimination tasks controlling for lesion size was also significant (*r*(72) = .0.32, *p* = .005), suggesting that overall lesion size is not accounting for the relationship between the two tasks.

**Figure 6:**
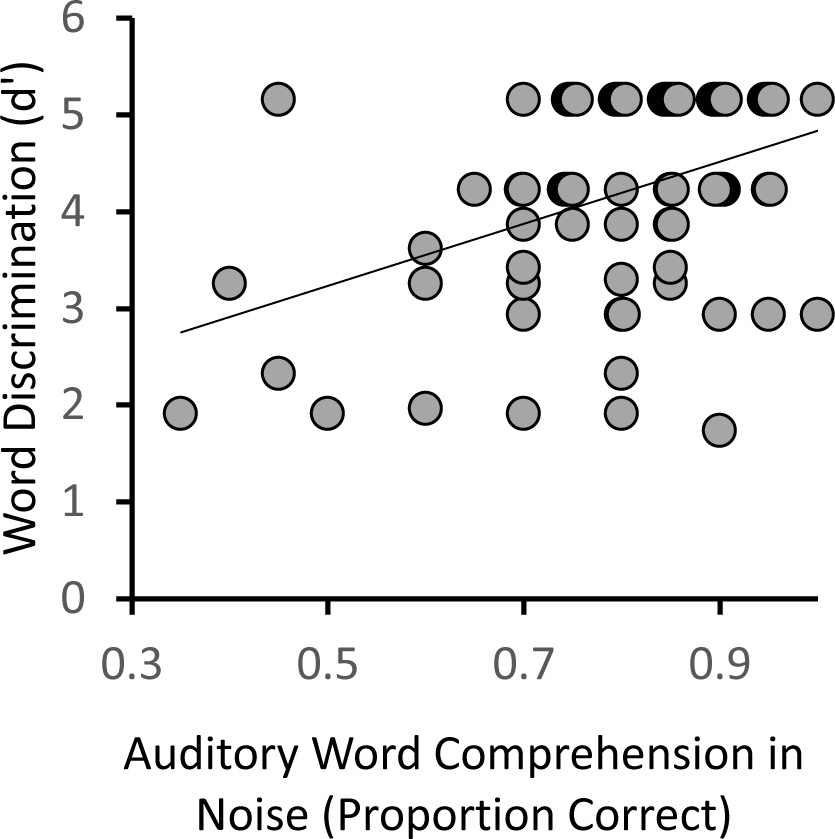
Relation between the auditory word comprehension in noise and word discrimination tasks within the LH stroke group. Overlapping points have been offset slightly to make them visible.

### Lesion-behavior mapping results

Lesion-symptom mappings (LSM) were computed within the left hemisphere participants for auditory word comprehension in noise, word discrimination, and nonword discrimination. The auditory word comprehension in noise task identified four adjacent ROIs covering much of the superior temporal lobe (Figure 7, Table 4). These ROIs included the superior and middle temporal gyri and underlying white matter, as well as extending into the inferior portion of the supramarginal gyrus; Heschl’s gyrus also was implicated as it is part of the anterior superior temporal ROI (JHU atlas label = “superior temporal gyrus,” *z* = 3.08, ROI center of mass = MNI coordinates -51 -13 1) (Figure 8A). The LSM of word discrimination identified two significant ROIs (*p*< .05, corrected; Table 4) adjacent to one another (JHU atlas labels “superior temporal gyrus” and “posterior temporal gyrus” Figure 7, 8B; Table 4), implicating most of the STG, as well as Heschl’s gyrus and the planum temporale, as well as an additional significant ROI, consisting of a portion of the superior longitudinal fasciculus (SLF) underlying the supramarginal gyrus (*z* = 3.46; ROI center of mass = -35 -24 29 MNI). The LSM of nonword discrimination identified the three ROIs implicated in real word discrimination (i.e., the two superior temporal gyrus ROIs, and the SLF ROI), as well as an additional ROI in the supramarginal gyrus extending into the post-central gyrus (*z* = 2.87, ROI center of mass = -51 - 29 33 MNI), which is dorsal to the supramarginal gyrus region implicated in the auditory word comprehension in noise LSM (Figure 7, 8B). All three LSMs overlap in the superior temporal gyrus ROI that includes mid-temporal regions including Heschl’s gyrus, and extends into the more posterior superior temporal gyrus (orange, Figure 8C; Table 4).

**Figure 7.**
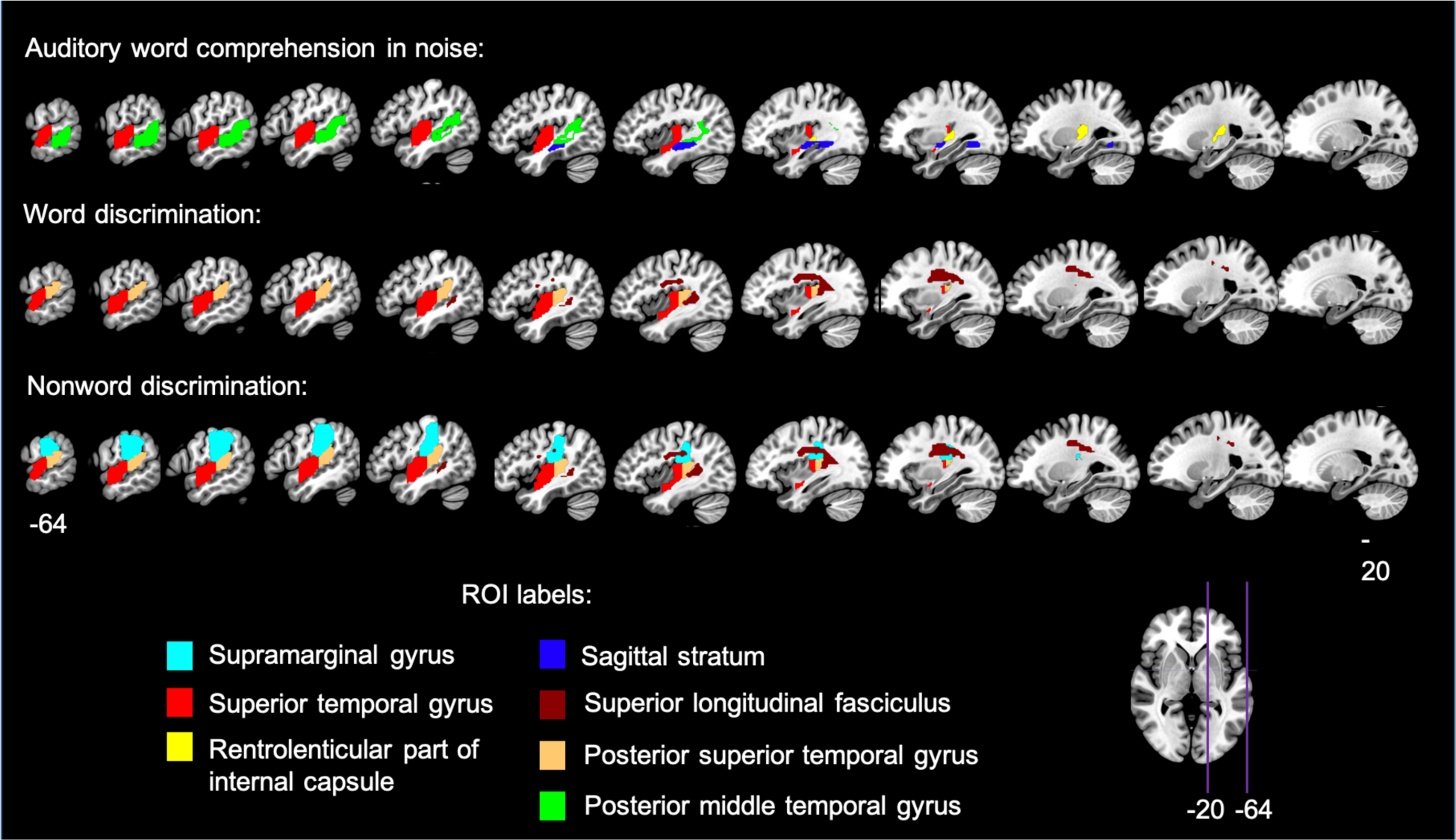
Lesion-symptom mapping results in the left hemisphere stroke group. Significant ROIs are displayed (p < .05, permutation-based family-wise error corrected) on sagittal slices through the left hemisphere for each of the three tasks for which lesion-symptom mapping was computed.

**Figure 8.**
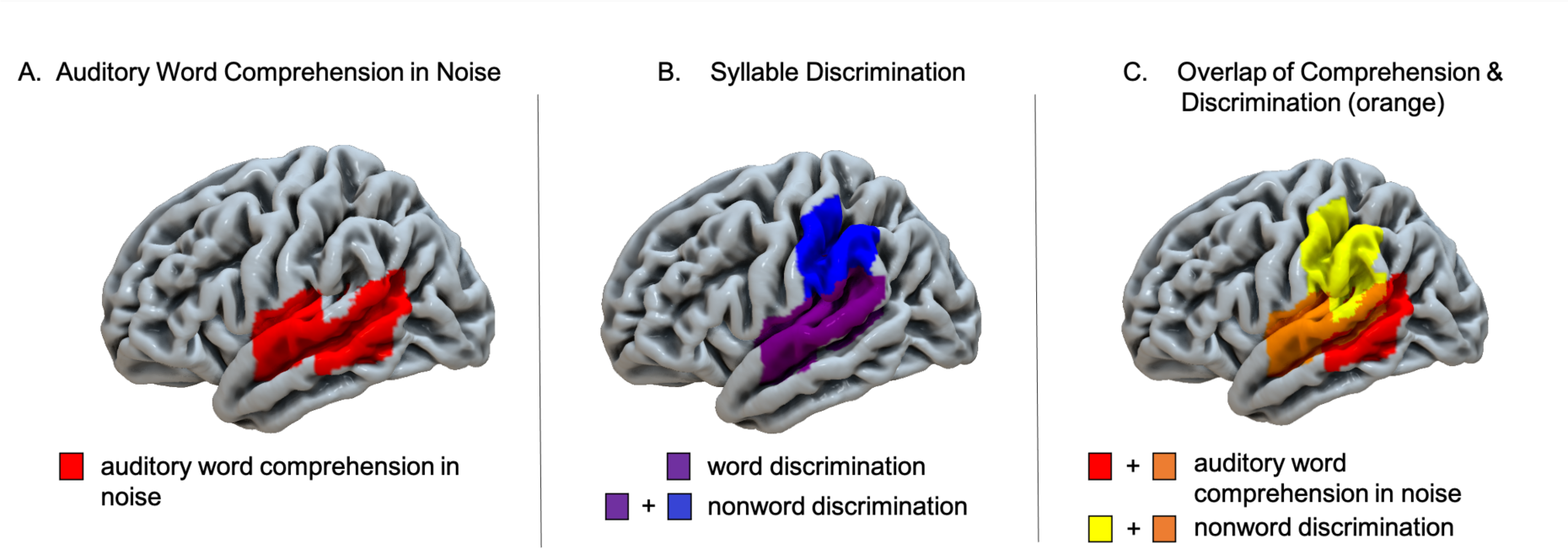
Lateral view summary of brain regions implicated in each task in the left hemisphere stroke group, p < .05, permutation-based family-wise error corrected. (A) Left lateral view of the regions significantly implicated in auditory word comprehension in noise (B) Left lateral view of the regions significantly implicated in word and nonword discrimination. Note that the word discrimination regions are a subset of the regions implicated in nonword discrimination. (C) Overlap of auditory word comprehension in noise and nonword discrimination indicated in orange.

**Table 4.**
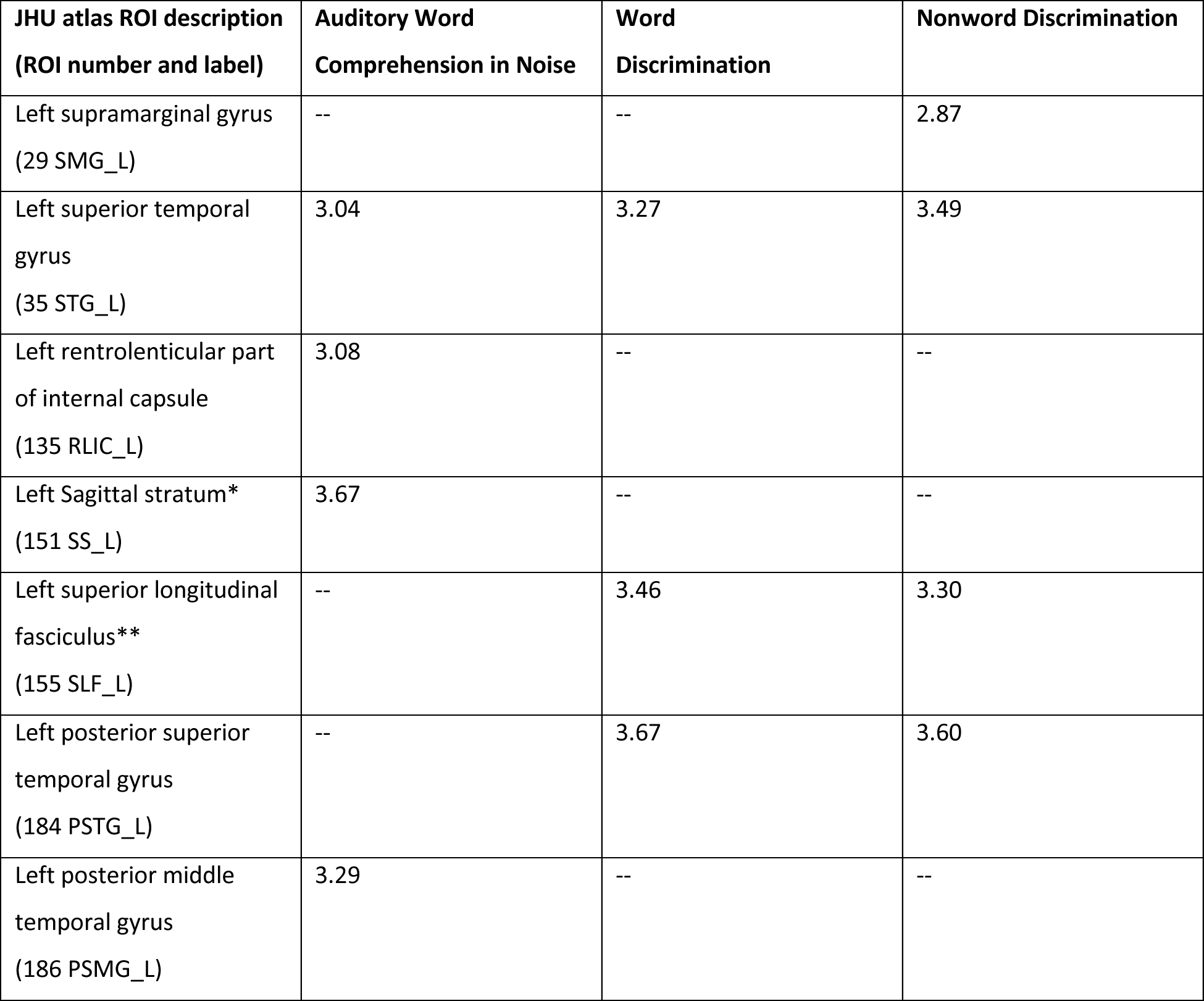
Z-scores of each significant ROI in the auditory word comprehension in noise task and the syllable discrimination tasks (i.e. word discrimination and nonword discrimination). *the JHU ROI atlas notes that the sagittal stratum includes inferior longitudinal fasciculus and inferior fronto-occipital fasciculus. **the JHU ROI atlas denotes this ROI as being part of the superior longitudinal fasciculus (SLF), but it is noteworthy that it is difficult to distinguish between SLF and arcuate fasciculus fibers in this region.

The results for the post-hoc conjunction and independence tests of the LSMs are as follows: for the word discrimination versus the nonword discrimination contrast (LSMs shown in Figures 7 and 8B), the conjunction test identified that damage in the left superior temporal gyrus (*z* = 3.12), the left posterior superior temporal gyrus (*z* = 3.04), and the left superior longitudinal fasciculus (*z* = 2.99) ROIs (i.e. all three ROIs that were separately significant for each task (Table 4)) were each related to significantly lower performance in both tasks (*p*< .05); the independence test did not identify any ROIs as being significantly more implicated in the nonword than the word discrimination task, or vice versa (i.e. *z* < 2.96, *p* > .05 in all ROIs). For the nonword discrimination versus auditory word comprehension in noise LSMs (LSMs shown in Figure 8C), the conjunction test identified damage in the left superior temporal gyrus ROI (*z* = 3. 21) as being significantly related to lower performance in both tasks (*p*< .05); the independence test identified damage in the left supramarginal gyrus (*z* = 3.21), left posterior superior temporal gyrus (*z* = 3.20), and left superior longitudinal fasciculus ROIs (*z* = 3.57) as being significantly more related to lower performance for nonword discrimination than for auditory comprehension in noise, and damage in the posterior middle temporal gyrus (*z* = 3.84) and sagittal striatum (*z* = 3.91) ROIs as being significantly more related to lower performance for auditory comprehension in noise than for nonword discrimination (*p* < .05). In short, the findings from the post hoc conjunction and independence analysis statistically confirm the map shown in Figure 8C.

### Right Hemisphere Results

To test for effects of right hemisphere damage on speech perception, we partitioned the RH Stroke group into those cases with (“Temporal +”) or without (“Temporal – “) damage (Figure 2 A,B) to the posterior superior temporal lobe and compared them to controls on the auditory word comprehension in noise task and on each syllable discrimination task. The one-way ANOVA across the three groups (RH Temporal +, RH Temporal -, and control) on the auditory word comprehension in noise task yielded a main effect of group that did not pass the *α* > .05 threshold but approached significance (F(2, 40) = 3.03, p = .061), with the superior temporal lobe damaged group performing numerically worse than those without damage there (Figure 8, left panel). A mixed ANOVA on the two syllable discrimination tasks across the same three groups revealed a significant effect of Task (F(1, 38) = 7.4, p = .01) with nonword discrimination performance lower than word discrimination, and a significant effect of Group (F(2, 38) = 7.5, p = .002), with the RH Temporal + group performing significantly worse than the controls and the RH Temporal- group (*p*s<.002, Bonferroni correction of *α* = .05/3 =.017); there was no significant difference between the control group and the RH Temporal- group (Figure 9, bottom). The Task x Group interaction was not significant (F(2,38) = 1.5, p=.24).

**Figure 9.**
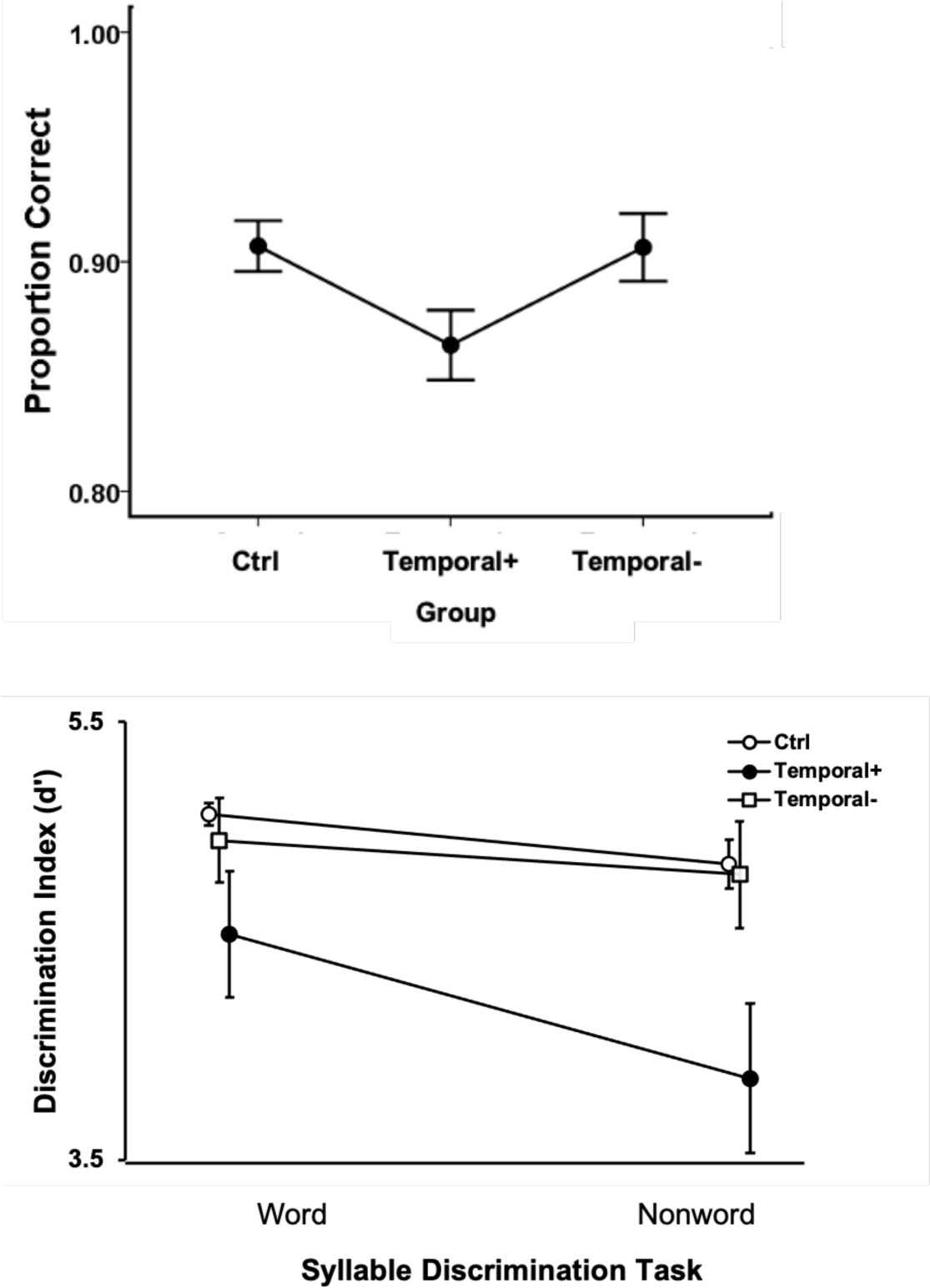
Right hemisphere auditory word comprehension in noise (top) and syllable discrimination performance (bottom). Error bars represent standard error. Overlapping points in the syllable discrimination graph (bottom) have been offset slightly to make them visible.

## Discussion

The present study sought to test four functional anatomic models of speech perception:

1. Classical model—speech perception is uniquely supported by the left mid-to-posterior superior temporal lobe.
2. Motor model—speech perception is dependent on (strong version) or augmented by (weak version) frontal motor systems involved in speech production, particularly under noisy listening conditions.
3. Left anterior temporal model—speech perception is dependent on the left anterior superior temporal lobe.
4. Bilateral temporal model—speech perception is bilaterally organized in the mid-to-posterior superior temporal lobe.

We also sought to evaluate potential behavioral and neural differences on two tasks that are often used to assess speech perception:

5. Task dependent claim—discrimination and comprehension tasks are dissociable both behaviorally and neurally.

To this end, we examined syllable discrimination and auditory word comprehension in chronic stroke, anterior temporal lobectomy, and unimpaired control participants. Consistent with prior work, we found that performance on auditory word comprehension and syllable discrimination tasks are dissociable behaviorally^2^. Performance on the clear auditory word comprehension task was at ceiling for the vast majority of the left hemisphere stroke group, precluding lesion analysis on that task. Lesion-symptom mapping using the auditory word comprehension in noise task as well as both syllable discrimination tasks revealed an association between each of these tasks and the left mid-to-posterior superior temporal gyrus. Task differences were also observed, however. The syllable discrimination tasks also uniquely implicated regions in the inferior parietal lobe and planum temporale, while the auditory word comprehension in noise task uniquely implicated a more ventral region in the middle temporal gyrus. Lesions involving the anterior temporal lobe or frontal, motor-related regions were not associated with poorer performance on any of the tasks, although the superior longitudinal fasciculus, which terminates in several areas of the frontal cortex including premotor regions, was implicated in both syllable discrimination tasks. Individuals with right hemisphere temporal lobe damage exhibited significant deficits on the syllable discrimination tasks and approached significance on the auditory word comprehension in noise task. In the following sections we discuss the implications of these findings for the specific hypotheses we aimed to test, starting with the task effects.

### Task dependence

Noting early reports of the paradoxical dissociability of auditory word comprehension tasks from syllable discrimination tasks (Basso, Casati, & Vignolo, 1977; S.E. Blumstein, Baker, & Goodglass, 1977; Miceli et al., 1980), Hickok and Poeppel argued that the two tasks involve shared resources in the superior temporal gyrus including what they termed “spectrotemporal analysis” in and around primary auditory cortex and extending into the ventral extent of the gyrus in the superior temporal sulcus, but then tapped into divergent processing streams beyond that (Hickok & Poeppel, 2007). They argued that discrimination tasks, in addition, relied on working memory systems in the motor-related dorsal stream (inferior parietal lobe, posterior planum temporale, and posterior frontal cortex) (Hickok et al. 2003; Buchsbaum & D’Esposito, 2019; Buchsbaum et al. 2011; Schomers et al. 2017; McGettigan et al. 2011), whereas auditory word comprehension tasks also relied on more ventral lexical access systems in the middle temporal gyrus region (Indefrey, 2011; Indefrey & Levelt, 2004; Lau, Phillips, & Poeppel, 2008; Liebenthal, Binder, Spitzer, Possing, & Medler, 2005; Pillay, Binder, Humphries, Gross, & Book, 2017).

Our findings are highly consistent with this hypothesis. Our study replicates previous work demonstrating the dissociability of syllable discrimination and auditory word comprehension tasks and identifies both shared and distinct regions implicated in the two tasks. The shared region involves the mid-to-posterior superior temporal gyrus, which was implicated in both word and nonword discrimination tasks, as well as the auditory word comprehension in noise task. The two discrimination tasks, but not the comprehension in noise task, additionally implicated the planum temporal region (part of the posterior superior temporal gyrus ROI), which includes area Spt, thought to serve as an auditory-motor interface for speech production (Buchsbaum et al., 2011; Hickok, 2012; Hickok, Buchsbaum, Humphries, & Muftuler, 2003; Hickok, Okada, & Serences, 2009). The nonword discrimination task uniquely implicated additional inferior parietal cortex in the supramarginal gyrus. In contrast, the auditory word comprehension in noise task, but not the discrimination tasks, implicated the posterior middle temporal gyrus, thought to be important for lexical-semantic access (de Zubicaray, Wilson, McMahon, & Muthiah, 2001; Hickok & Poeppel, 2004, 2007; Indefrey, 2011; Indefrey & Levelt, 2004; Lau et al., 2008).

Given that functional imaging and TMS have implicated frontal, motor-related regions in the performance of syllable discrimination and related tasks (S. E. Blumstein, Myers, & Rissman, 2005; Burton et al., 2000; D’Ausilio et al., 2009; Jancke, Wustenberg, Scheich, & Heinze, 2002; Lee, Turkeltaub, Granger, & Raizada, 2012; Meister et al., 2007; Watkins & Paus, 2004; Zatorre, Evans, Meyer, & Gjedde, 1992), we expected to see some frontal ROIs correlated with deficits on our discrimination task. Yet we did not. This result does not necessarily mean that motor speech systems play no causal role at all in speech processing using some tasks and under some conditions. The TMS studies suggest otherwise as does the present finding that temporal-parietal dorsal stream regions that play an important role is speech production are implicated in syllable discrimination. Presumably, these temporal-parietal areas are interfering with discrimination task performance as a function of their network connectivity with frontal, motor-related areas. Our finding of damage to the superior longitudinal fasciculus being associated with poorer performance on both syllable discrimination tasks, but not the comprehension task, also supports view. Yet for reasons that are not entirely clear, disrupting posterior nodes in the dorsal stream sensorimotor network seems to have the dominant effect on syllable discrimination relative to damage in frontal motor speech areas (Caplan, Gow, & Makris, 1995).

What is clear, is that none of these dorsal stream regions, when damaged, impair the ability to comprehend words. Further additional support for this conclusion comes from a recent study of two cases of acquired opercular syndrome, both with complete anarthria secondary to bilateral removal of most of the inferior, posterior frontal cortex. Despite being completely unable to voluntarily control their vocal tracts, both individuals were able to comprehend words at ceiling levels (Walker, Rollo, Tandon, & Hickok, 2021).

Our neuroanatomical findings for these tasks explain the behavioral patterns of association and dissociation. Overall, across the LH stroke group, word discrimination and word comprehension (in noise) are weakly but significantly correlated (Figure 6). This co-variation is presumably related to shared involvement of auditory and phonological systems in the superior temporal gyrus. The dissociability of the two tasks (Figure 6 and Table 3) is explained on the basis of the distinct pathways recruited by the two tasks beyond the superior temporal gyrus. Damage to dorsal stream systems can impact discrimination without affecting comprehension and damage to ventral temporal lobe systems can impact comprehension without affecting discrimination.

### The classical model of speech perception

The classical model of speech perception, originally proposed by Wernicke, was not supported by the present data. This model, which holds that the left superior temporal gyrus is the primary substrate for speech perception, predicts that damage to this region should cause substantial deficits on all receptive speech tasks. This was not the case as the vast majority of individuals in our LH stroke group were at ceiling on the clear auditory word comprehension task even though the task required the ability to resolve minimal phonemic pair differences. This finding not only shows that damage to classical Wernicke’s area is not sufficient for producing substantial deficits in word comprehension, but more generally that damage anywhere in the left hemisphere is not sufficient. This, in turn, indicates that receptive speech up to the word level is not strongly lateralized and is more consistent with a bilateral model, which we discuss below.

### The motor models of speech perception

Our study provided little support for the strong motor model, which predicts that damage to motor speech areas should cause deficits in the ability to perceive speech sounds and therefore affect both syllable discrimination and auditory word comprehension. As noted, clear auditory word comprehension was largely unaffected by unilateral stroke even with a substantial number of individuals in the LH stroke group exhibiting clinically significant motor speech deficits (e.g. Broca’s aphasia). Further, damage to frontal motor-speech areas was not significantly correlated with deficits on any of our tasks. As pointed out above, the motor model does gain partial support in that damage to temporal-parietal dorsal stream regions, which are part of the speech production network, are implicated in performance on discrimination tasks. However, these effects do not generalize to the more ecologically valid comprehension tasks.

Weaker versions of the motor model also did not gain support from our study. These models argue that motor speech systems play an augmentative role in speech perception under noisy listening conditions (Moineau et al., 2005; Wilson, 2009). We studied auditory word comprehension in noise and found that damage to superior temporal and middle temporal regions was correlated with deficits; dorsal stream regions were not implicated. This finding contradicts a previous, smaller-scale study that reported that word comprehension suffered with degraded speech in people with Broca’s aphasia, who typically have damage to frontal, motor-related speech areas (Moineau et al., 2005). There are two difficulties interpreting this previous study, however. One is that Broca’s aphasia is not caused solely by damage to frontal cortex and indeed regularly also involves damage to classical Wernicke’s area in the mid and posterior temporal lobe (Fridriksson et al., 2015), i.e. regions that our study linked to deficits on the auditory word comprehension in noise task. Thus, it could be that it was damage to Wernicke’s area rather than motor speech areas that was the source of the effect reported by Moineau et al. The second difficulty is that the behavioral analysis did not take response bias into account. The task was to decide whether a word-picture pair matched or did not. Performance on such a task is affected not only by perceptual ability but also response bias. Previous work using fMRI (Venezia et al., 2012) and lesion (Rogalsky et al., 2018) methods have linked motor speech areas to changes in response bias. Moineau et al. report significant response bias effects in their data, yet report only percent correct measures rather than d-prime statistics, which would correct for bias (Green & Swets, 1966). Thus, even if the lesion distribution in Broca’s aphasia was not a confound, it is unclear if Moineau et al.’s observed effects are due to perceptual effects or changes in response bias. The data from the present, larger-scale study that includes a direct analysis of lesion patterns suggest that one or both of the confounds in the Moineau et al. study impacted their findings.

One could argue that our study did not include the full range of tasks that have been shown to implicate frontal motor-related systems in speech processing. For example, Burton et al. (2000) showed that discriminating the onset phoneme in syllable pairs such as *dip-tip* did not consistently yield frontal activation whereas in pairs such as *dip-ten* it did, which they argued was due to explicit phoneme segmentation (similar to a phonological awareness task) or working memory. Our task used stimuli more similar to *dip-tip,* leaving open the possibility that a different task requiring explicit segmentation or working memory may recruit frontal motor-related systems. But such a possibility does not weaken our claim that speech perception as it is employed under more natural conditions does not substantively involve the frontal motor speech system. Rather, it would strengthen the view that motor involvement is an artifact of tasks that require motor-related processes, such as working memory (Hickok & Poeppel, 2000, 2004, 2007).

### The anterior temporal model of speech perception

The anterior temporal model predicts that damage to the left anterior temporal lobe should have a substantial impact on the ability to comprehend speech. Our findings provide partial support for this model in that, for all tasks, lesions implicated mid-posterior as well as mid-anterior STG regions. The “left superior temporal gyrus” ROI that was implicated in all tasks includes mid-anterior STG and extends anteriorly such that it nearly included the coordinates for fMRI activations in the ATL that have been claimed to be the foci involved in intelligible speech processing (Figure 10), as revealed in studies of intelligible versus unintelligible sentences (Scott et al. 2000; Narain et al. 2003; Evans et al. 2014).

**Figure 10.**
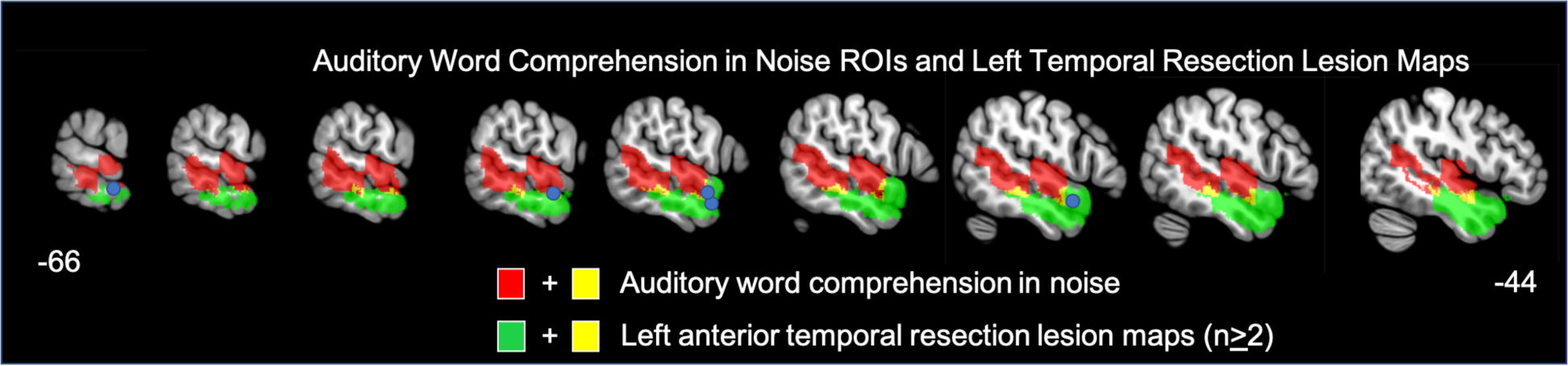
Red: Significant ROIs identified by the LSM analyses for the auditory word comprehension in noise task; Green: The left lobectomy group’s lesion maps (areas shown indicate lesion in at least 2 participants); Yellow: overlap between the auditory word comprehension in noise ROIs and the left lobectomy lesion maps. Blue circles indicate the five anterior temporal coordinates identified by Scott et al. 2000 (MNI: -54 6 -16, -66 -12 -12) or Evans et al. 2014 (MNI: -58 2 -16, -60 -2 -12, -58 4 -20) as exhibiting main effects of speech intelligibility or a similar contrast.

These fMRI activation foci largely fall within the region impacted by the present study’s left hemisphere ATL resection cases, which as a group scored lower than controls and similar to the group average for individuals with right posterior temporal strokes, but quite a bit better than the left hemisphere stroke group. This suggests a role for the left ATL in speech perception, but does not support the claim that the network is strongly left dominant (Scott et al. 2000; (Rauschecker & Scott, 2009) as more severe deficits should have been apparent with left hemisphere damage, even on the clear auditory word comprehension task if the network was strongly left dominant. Further, in the context of the anterior temporal lobe model, it has been claimed that posterior superior temporal regions primarily support auditory-motor functions, not intelligibility (Rauschecker & Scott, 2009) thus predicting that posterior temporal damage should not be implicated in word comprehension. Our findings suggest otherwise and argue that the “pathway for intelligible speech” involves much of the superior temporal gyrus and extends into the posterior middle temporal gyrus consistent with higher-powered functional imaging studies on intelligible speech (Evans et al. 2014; Okada et al. 2010).

### The bilateral temporal model of speech perception

The bilateral temporal model predicts that speech perception as measured by auditory word comprehension will not be substantially impaired by unilateral damage to either hemisphere and that any mild deficits will be associated with superior temporal lobe damage. This is precisely the result we found. Clear speech word comprehension was at or near ceiling in the vast majority of all people studied and poorer performance on the auditory word comprehension in noise task was associated with damage to the superior temporal gyrus and posterior middle temporal gyrus. This was true in the left hemisphere where we had a sufficient sample size to perform lesion-symptom mapping; a non-significant trend (p = .06) in the same direction was found as well in the 11 participants with right hemisphere mid-posterior superior temporal lobe lesions.

Previous authors have argued for a bilateral capacity for receptive speech at the word level based on data from split brain studies, Wada procedures, and stroke (Hickok et al., 2008) Gazzaniga & Sperry, 1967; Zaidel, 1985; Buchman et al. 1986; Bachman & Albert, 1988; Goodglass, 1993; Hickok & Poeppel 2000, 2004, 2007). Until recently, however, direct evidence has been lacking for an association between speech perception impairment and unilateral right hemisphere damage, although higher-level language deficits have been documented (Gajardo-Vidal et al., 2018). The present study provides such evidence from stroke and joins a recent TMS study reporting similar effects when left or right superior temporal regions were interrupted in healthy participants (Kennedy-Higgins et al. 2020).

A recent large-scale lesion study using a clinical assessment tool (WAB-R) reported that unilateral left hemisphere stroke can cause single word comprehension deficits (in clear speech), which map onto left temporal lobe regions (Bonilha et al., 2017). However, scores on this task include subtests that assess aspects of reading (matching spoken words to printed letters and numbers) as well as shapes, colors, and body parts in addition to more common objects. In light of the high level of performance on the present study’s clear auditory word comprehension task, it is likely that clinical assessment tools tap into a broader range of abilities that go beyond basic phonemic perception and lexical-semantic access, which is what our tasks were designed to test.

Although group performance was exceptionally high on the clear auditory word comprehension task (mean = 97%) with 95.4% of our sample performing at or better than 90% correct, there were a handful of cases (∼5%) who were substantially impaired. How should these cases be interpreted with respect to claims about the lateralization pattern for auditory word comprehension? We suggest that these cases likely reflect anomalous dominance, that is, cases where auditory word comprehension is atypically dependent on the left hemisphere. In general, the incidence of anomalous cerebral lateralization for language in strong right handers is in that same range with estimates at approximately 5% (Knecht, et al. 2000) and functional brain imaging work suggests the incidence of almost complete left dominance is itself atypical and with a frequency of approximately 5% (Springer, et al. 1999). One might counter that because aphasia was not an inclusion criterion at all of our testing sites, the actual incidence of word comprehension impairments following a left hemisphere stroke might be underestimated in our sample. Yet even at sites that recruited only people with aphasia, only 4 of 63 (6.3%) performed worse than 90% correct on the clear word comprehension test. Moreover, the reverse sampling bias argument can be made: recruiting participants on the basis of the presence of aphasia following unilateral stroke likely *over estimates* left hemisphere dominance for language because people with naturally more bilateral organization will be less likely to present with aphasia following unilateral stroke.

A final potential concern regarding the present data for the bilateral model is whether plastic reorganization is responsible for the well-preserved word comprehension ability. That is, perhaps performance is at ceiling because the right hemisphere is taking on speech recognition function after left hemisphere insult. However, data from a large sample of acute stroke (Rogalsky et al. 2008) as well as a smaller sample of acute left hemisphere deactivation in a Wada study (Hickok et al., 2008) show similar effects.

Although our findings strongly support the bilateral model, this does not necessarily imply symmetry of function (Zatorre *et al*. 2002; Poeppel, 2003; Hickok & Poeppel, 2007; Morillon et al. 2010). Indeed, we found greater impairment following left compared to right hemisphere damage on our auditory word comprehension in noise task as well as our discrimination tasks. This could reflect left-right computational differences within the speech perception systems (Poeppel, 2001; Zatorre et al. 2002; (Albouy et al., 2020) or their interaction with other more lateralized networks—lexical-semantic, sensorimotor, attention—that may augment the efficiency of the left hemisphere for processing speech for some tasks.

### Summary

In summary, the findings from the present study support a neuroanatomical model of basic auditory word identification and comprehension in which the critical regions include much of the superior temporal lobe, bilaterally: a hybrid of the anterior temporal and bilateral models discussed above. This work also confirmed the dissociability, both behaviorally and anatomically, of comprehension- and discrimination-based auditory speech tasks. Given that a large of number of studies of “speech perception” use discrimination-type tasks, particularly those that implicate dorsal, motor-related networks (D’Ausilio et al. 2009; Meister et al. 2007; Mottonen et al. 2009), the present finding calls for a critical re-evaluation of that body of work.

## Footnotes

1 The participants with temporal lobectomies have lesions localized in the anterior temporal lobe (Fig 2), a region that is often left intact by all but the most severe middle cerebral artery strokes (Holland & Lambdon Ralph, 2010) and therefore of particular value to the present study. However, there is evidence that epileptic brains have variable functional organization, including laterality of speech processing (Swanson et al. 2002, 2007; Hertz-Pannier et al. 2002). Thus, findings in this population should be interpreted carefully.

2 One possible explanation of these tasks being dissociable might be general task difficulty differences, but there are two strong pieces of evidence against a general task difficulty effect and indicating a speech perception-related effect: (1) the ROIs identified are in auditory-related areas and not frontal regions typically implicated in task demands, and (2) the vast majority of the errors induced by adding noise were phonemic errors (i.e. selecting the phonemic foil picture).

